# FSL-MRS: An end-to-end spectroscopy analysis package

**DOI:** 10.1101/2020.06.16.155291

**Authors:** William T Clarke, Charlotte J Stagg, Saad Jbabdi

**Affiliations:** Wellcome Centre for Integrative Neuroimaging, FMRIB, Nuffield Department of Clinical Neurosciences, University of Oxford, Oxford, United Kingdom

**Author notes:** To whom correspondence should be addressed **Corresponding author contact details:** Dr William T. Clarke, Wellcome Centre for Integrative Neuroimaging, FMRIB, University of Oxford, Level 0, John Radcliffe Hospital, Oxford, OX3 9DU, UK.

**Keywords:** Spectroscopy, MRS, MRSI, Bayesian fitting, open-source

## Abstract

**Purpose:** We introduce FSL-MRS, an end-to-end, modular, open-source magnetic resonance spectroscopy analysis toolbox. FSL-MRS provides spectroscopic data conversion, pre-processing, spectral simulation, fitting, quantitation and visualisation.

**Methods:** FSL-MRS is modular. FSL-MRS programs operate on data in a standard format (NIfTI) capable of storing single voxel and multi-voxel spectroscopy, including spatial orientation information.

FSL-MRS includes tools for pre-processing of raw spectroscopy data, including coil-combination, frequency and phase alignment, and filtering. A density matrix simulation program is supplied for generation of basis spectra from simple text-based descriptions of pulse sequences.

Fitting is based on linear combination of basis spectra and implements Markov chain Monte Carlo optimisation for the estimation of the full posterior distribution of metabolite concentrations. Validation of the fitting is carried out on independently created simulated data, phantom data, and three in vivo human datasets (257 SVS and 8 MRSI datasets) at 3T and 7T.

Interactive HTML reports are automatically generated by processing and fitting stages of the toolbox. FSL-MRS can be used on the command line or interactively in the Python language.

**Results:** Validation of the fitting shows low error in simulation (median error 11.9%) and in phantom (3.4%). Average correlation between a third-party toolbox (LCModel) and FSL-MRS was high (0.53-0.81) in all three in vivo datasets.

**Conclusion:** FSL-MRS is designed to be flexible and extensible to new forms of spectroscopic acquisitions. Custom fitting models can be specified within the framework for dynamic or multi-voxel spectroscopy. FSL-MRS is available as part of the FMRIB Software Library.

## Introduction

Recent years have seen the emergence and rapid progress of new magnetic resonance spectroscopy (MRS) technologies, including spectral editing (1), MRS imaging (2,3), time-resolved functional MRS (4), diffusion-weighted MRS (5), and MRS fingerprinting (6). MRS is therefore starting to have a range of techniques comparable to those of conventional proton MRI, but with the added benefit of being able to quantify specific chemical compounds. However, unlike modern MRI-based neuroimaging, MRS lacks standard data formats (e.g. NIfTI (7)), as well as standard pre-processing and analysis pipelines suitable for use by non-expert users (e.g. FSL (8), SPM (9), or AFNI (10)). This restricts the use of MRS in research, particularly in neuroscience, by requiring expertise in MRS acquisition, data analysis, and computing. Current processing toolboxes are typically linear and linearly dependent, lacking modularity or a standardised data format. It is therefore difficult to customise processing pipelines, inspect the results of single steps of a pipeline, or combine steps from different toolsets.

The tools currently available and commonly in use for processing, fitting and visualisation of spectra (e.g. (11-16)) suffer from one or more of several limitations, namely:

- They may be black-box, closed-source implementations, sometimes with monetary cost.
- They may require licenced software to run, which is not universally deployable.
- They often require high user interaction, either through a GUI or the need for setting and understanding many options.
- They have fixed forward fitting models. Modifications require MRS and computing expertise.
- They have limited or no handling of MRSI data, with no parallelisation available.

For these reasons currently available software is not easily extensible to new forms of MRS, for example high-resolution, high-voxel count MRSI, or time-series modelling of functional MRS (fMRS), or diffusion-weighted MRS (dwMRS).

In this work we present a new Python-based MRS fitting and processing tool, FSL-MRS. The toolbox is open-source, free as part of the FSL software package (8), and operates with a scriptable command line or interactive interface. It implements a modular approach to spectroscopy analysis with a common data format, allowing integration with other neuroimaging tools. Steps are parallelisable for MRSI data. FSL-MRS is end-to-end, comprising modules for data conversion, pre-processing, basis spectra simulation, fitting, quantification, and visualisation.

The FSL-MRS fitting module works on the principle of linear combination of pre-calculated basis spectra (13). In keeping with FSL’s tradition of favouring Bayesian inference approaches (17), our tool calculates full posterior distributions of the fitted metabolite concentrations using a Markov chain Monte Carlo (MCMC) algorithm, specifically Metropolis-Hastings (18). The full posterior distributions can be utilised in further analysis, allowing efficient propagation of fitting uncertainties into downstream modelling and statistical analyses. Parameter covariances are also available from the fitting output, and point estimates of concentration and uncertainties may be calculated using appropriate summary statistics. FSL-MRS incorporates an interactive reporting interface which uses modern data-science visualisation tools.

In this work we describe the FSL-MRS components, interface and output, and describe the fitting model and approach. A validation of the tool’s fitting estimates is carried out on widely available simulated data, in phantom and on three in vivo datasets at 3T and 7T, spanning 265 subjects.

## Methods

### Data conversion and format

FSL-MRS operates on a modular processing principle. Modularity allows custom third-party additions to the processing pipeline without the need to alter the FSL-MRS package or adhere to FSL-MRS imposed code conventions, languages, or possible limitations.

To enable this workflow FSL-MRS processing and fitting operates on MRS data stored in the Neuroimaging Informatics Technology Initiative (NIfTI) format (7). The NIfTI format permits the storage of data resolved into three spatial dimensions, in addition to a time dimension and two further unspecified dimensions. MRS and MRSI time-domain data may therefore be stored using the format (and it will also allow analysis of fMRS and dwMRS data in the future). Data is loaded from, and written to, file after each operation. Additional required meta-data is stored in, read from, and written to JavaScript Object Notation (JSON) “sidecar” files, as specified by the Brain Imaging Data Structure (BIDS) format (19).

FSL-MRS provides the spec2nii program to convert from existing data formats to NIfTI. Spec2nii currently supports seven formats specified in the supporting information (Supporting Information Table 1). Spectroscopy volume position information is translated into the NIfTI “qform” field where it is available in the original format.

**Table 1:** Processing operations available using the fsl_mrs_proc command line tool

| **fsl_mrs_proc operation** | **Description** | **Reference** |
| --- | --- | --- |
| coilcombine | Combine individual coils of receiver phased array. | (21,22) |
| average | Average FIDs, with optional complex weighting. |  |
| align | Phase and frequency align FIDs using spectral registration. | (23) |
| align-diff | Phase and frequency align sub-spectra based on addition or subtraction of sub-spectra (e.g. for ISIS localisation). | (23) |
| ecc | Eddy current correction using a water phase reference scan. | (25) |
| remove | Remove peak (typically residual water) using HLSVD. | (24) |
| tshift | shift/resample in time domain. |  |
| truncate | Truncate/pad time-domain data by an integer number of points. |  |
| apodize | Apply choice of apodization function to the data. |  |
| fshift | Frequency domain shift. |  |
| unlike | Identify outlier FIDs and remove based on similarity metric. | (14) |
| phase | Zero-order phase spectrum by phase of maximum point in range |  |
| subtract | Subtract two FIDs |  |
| add | Add two FIDs |  |

### Modular end-to-end processing

FSL-MRS provides a complete set of command line tools for spectroscopy analysis. Here we define *processing* as the steps required to make single voxel spectroscopy or reconstructed MRSI k-space data ready for *fitting. Basis spectra creation* is the process of using quantum mechanical simulations (or other methods) to create numerical descriptions of a metabolite’s response to a specific MRS pulse-sequence. *Fitting* is the process of estimating relative metabolite concentrations from the processed spectrum and the basis spectra. *Quantification* turns those relative concentrations into real-world interpretable units of concentration. *Display* incorporates viewing of the data and results at all stages of the process. Figure 1 shows an overview of the tool’s workflow.

**Figure 1 –.**
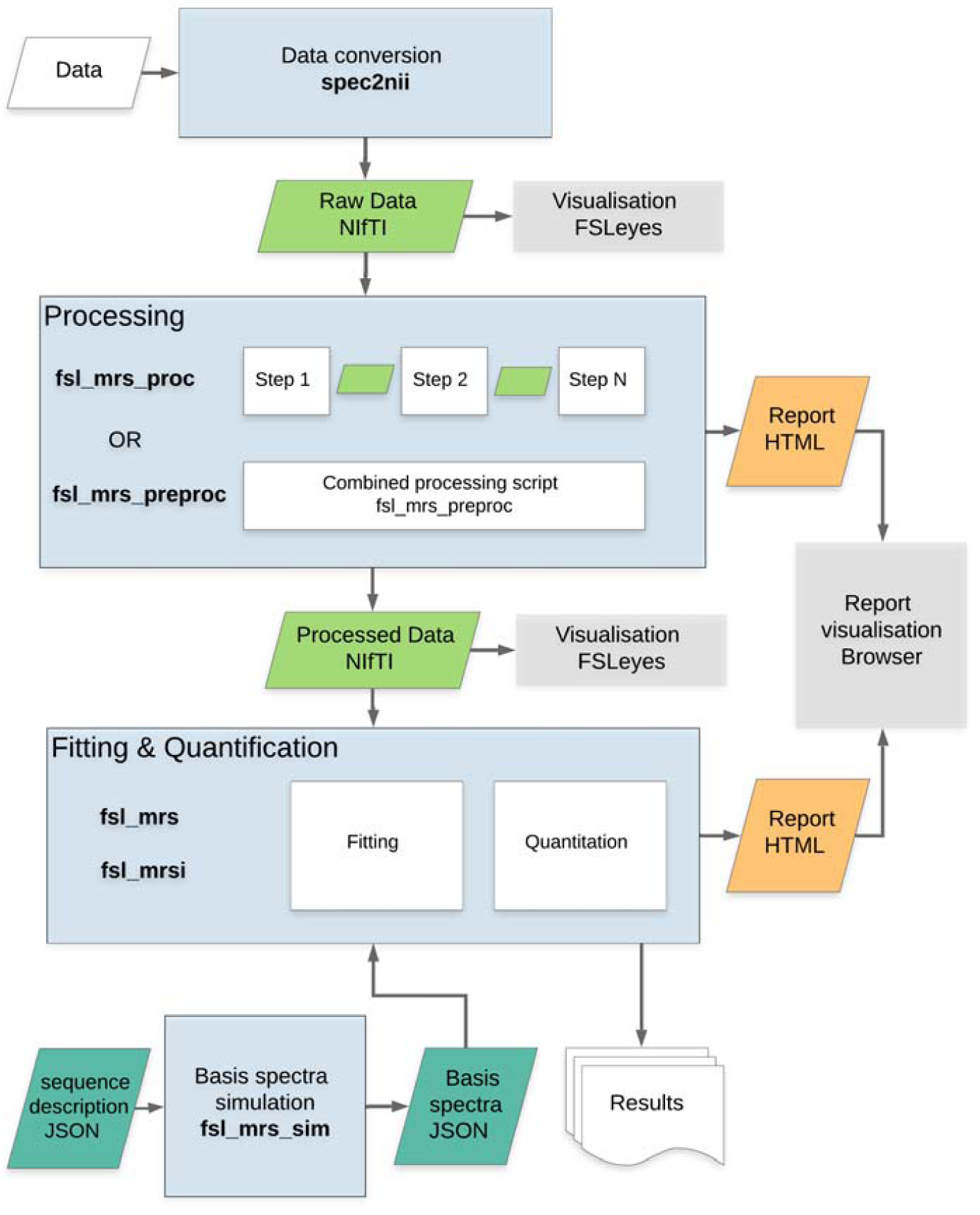
FSL-MRS organisation and workflow. Raw data in proprietary or other formats are converted to NIfTI by spec2nii. Processing can then be carried out in stages, operating on NIfTI files, using fsl_mrs_proc, or in a single python script fsl_mrs_preproc for standard SVS sequences. Basis spectra can be generated for fitting using fsl_mrs_sim given a json description for the sequence. Fitting and quantitation are then carried out by fsl_mrs and fsl_mrsi as appropriate. Interactive HTML reports are generated for viewing in the user’s browser. Spectroscopy data in NIfTI format can be viewed overlaid with other MR contrasts in FSLeyes.

### Processing

FSL-MRS provides tools for all the processing operations recommended in the community-driven consensus paper (tables 2, 3 and 4 of reference (20)). These tools are accessed through the command line by *fsl_mrs_proc* and are described briefly in Table 1. Coil combination is performed through the wSVD algorithm (21,22), spectral alignment by spectral registration (23), and nuisance peak removal by HLSVD (HLSVDPRO)(24). *fsl_mrs_proc* operations are applied sequentially to data stored in NIfTI format. Operations can be combined, in order, to form a repeatable batch processing script.

**Table 2.** Description of in vivo datasets used for validation. ACC = anterior cingulate cortex, OCC = occipital cortex, PCC = posterior cingulate cortex.

| # | Sequence | B<sub>0</sub> (T) | Subjects | Voxels (brain regions) | Measured MM? | Vendor | Ref |
| --- | --- | --- | --- | --- | --- | --- | --- |
| 1 | STEAM | 7 | 37 | 3 (ACC, OCC, putamen) | Yes | Siemens | (40) |
| 2 | SPECIAL | 3 | 220 | 1 (PCC) | Yes | Siemens | (42) |
| 3 | CONCEPT | 3 | 8 | 126 (calcarine sulcus) | No | Siemens | (43) |

**Table 3.** Summary of in vivo validation – correlation and Bland-Altman statistics. *All values are mean ± standard deviation. All = all metabolites (excluding combined), combined = after combination (excludes those combined), LoA = limits of agreement (width of 95% CI)*

| Dataset | Scaling | Correlation<br>*all* | Correlation<br>*combined* | % Bias<br>*all* | % Bias<br>*combined* | % LoA<br>*all* | % LoA<br>*combined* |
| --- | --- | --- | --- | --- | --- | --- | --- |
| 1) 7T<br>STEAM | Water<br>tCr | 0.69±0.18<br>0.71±0.17 | 0.74±0.19<br>0.75±0.18 | 22±21<br>20±21 | 14±12<br>11±11 | 120±71<br>108±63 | 93±64<br>84±64 |
| 2) 3T<br>SPECIAL | Water<br>tCr | 0.68±0.13<br>0.77±0.18 | 0.75±0.12<br>0.81±0.14 | 44±25<br>34±34 | 45±27<br>28±33 | 414±329<br>277±337 | 369±345<br>259±408 |
| 3) 3T<br>CONCEPT | Water<br>tCr | 0.53±0.16<br>0.54±0.15 | 0.56±0.14<br>0.58±0.13 | 37±22<br>27±22 | 37±20<br>24±23 | 216±109<br>162±90 | 193±92<br>138±96 |

In addition to the flexibility offered by this script, FSL-MRS also provides a pre-packaged processing pipeline for non-edited single voxel data – *fsl_mrs_preproc*, which runs all appropriate steps with one command-line operation.

### Basis spectra simulation

Fitting in FSL-MRS works on the principle of Linear Combination (LC) modelling (see Bayesian Fitting). LC modelling requires that the user provides the algorithm with simulated (or measured) numerical responses of metabolite spin systems to the MRS pulse sequence being used. These responses are specific to the pulse sequence, the sequence timings, and the sequence radio frequency (RF) pulse envelopes and are known as basis spectra. Basis spectra must preserve the relative signal amplitude between metabolites.

FSL-MRS provides an interface (*fsl_mrs_sim*) for the creation of basis spectra when provided with a description of the sequence timings, RF pulses, slice selection gradients and rephasing gradient areas. RF pulses may have arbitrary amplitude and phase modulation (i.e. be non-ideal). The description is provided in a JSON format, examples are provided in the software documentation. The simulation is based on the extended 1D projection implementation of density matrix simulations (26,27). Unwanted coherences are removed with a coherence order filter (28). Standard literature values for common spin system chemical shifts and coupling constants are included in the software (29,30).

*fsl_mrs_sim* outputs a JSON file for each simulated metabolite which may be loaded by FSL-MRS’s fitting modules. FSL-MRS also accepts LCModel (.BASIS) and jMRUI (.txt) basis spectra formats.

### Fitting & Quantification

Fitting in FSL-MRS is provided by two command-line interfaces: *fsl_mrs* (for single voxel spectroscopy [SVS] data) and *fsl_mrsi* (for MRSI data). Additional interfaces will be added in the future for other types of MRS (e.g. diffusion-weighted MRS, functional MRS). Fitting is carried out on each voxel of data independently. The user may optionally specify: the limits of fitting (in ppm), the order of the complex polynomial baseline (see Standard fitting model), whether to add default macromolecular peaks (at 0.9, 1.2, 1.4, 1.7 and 2.08 & 3.0 ppm), the optimisation algorithm (see Optimisation). Metabolites in the basis spectra file may be optionally excluded by the user and, for output purposes only, metabolites may be combined.

For meaningful quantification the user must supply a processed unsuppressed water dataset, and for transverse relaxation-compensated concentrations the user must supply the sequence echo time and tissue volume fractions (31,32). Water signal amplitude (*S*_H2O_obs_ in references (20,31,32)) is calculated using numerical integration of the real part of the phase corrected and eddy current corrected, unsuppressed water spectrum. Water-scaled concentrations are calculated by taking the ratio to the integrated signal of a scaled reference metabolite basis spectrum (*S*_M_obs_ in references (20,31,32), default of creatine between 2 and 5 ppm). The scripts *svs_segment* and *mrsi_segment* can calculate tissue volume fractions within each voxel given an appropriate T1-weighted structural image. Default values for water concentration, tissue-water densities, and water and metabolite T_2_ time constants are provided for 3T and 7T field strengths. Hardcoded constants, correct as of time of publication, may be found in the supporting information (Tables 2-4), and the values correct for the current version may always be found in the source-code module *fsl_mrs*.*utils*.*constants* or as part of the online documentation. These defaults may be overridden in the interactive or python interface. Concentrations can be expressed as a ratio to an arbitrary internal reference metabolite (or combination of metabolites) or in molar (mol/dm^3^) or molal (mol/kg) units.

FSL-MRS fitting outputs the signal-to-noise ratio (SNR) and linewidths (full width at half maximum) for each fitted metabolite. The SNR ratio is calculated as the ratio of the peak height of the fitted metabolite basis spectrum over the standard deviation of a pure noise region of the spectrum after a matched filter has been applied to both (33). The matched filter and linewidth are calculated for each metabolite as the FWHM peak width in hertz, as calculated from the most prominent peak in the fitted basis spectrum. If the MCMC algorithm is used the quality control metrics are calculated over all samples.

### Reporting & Display

FSL-MRS modules generate self-contained interactive HTML reports (Plotly, Montreal, Canada) which can be viewed and interacted with in the user’s web browser. All components of the processing module (Table 1) produce short HTML reports which can be combined into a single interactive report for each instance of data by using the packaged *merge_mrs_reports*.

An interactive report is generated for each fit displaying the fitted spectrum, model fit, residuals and concentrations (Fig 1) and concentration posterior distributions, metabolite covariances and scaled basis spectra (Fig 2). The report also contains summaries of SNR and linewidth quality control parameters for each fitted metabolite. The user therefore can quickly assess the quality of SVS data and fit visually in one location. Results of the fitting, and quality control metrics, are also available as comma-separated values (CSV) files from the command-line programs and as Pandas objects in memory (34).

**Figure 2 –.**
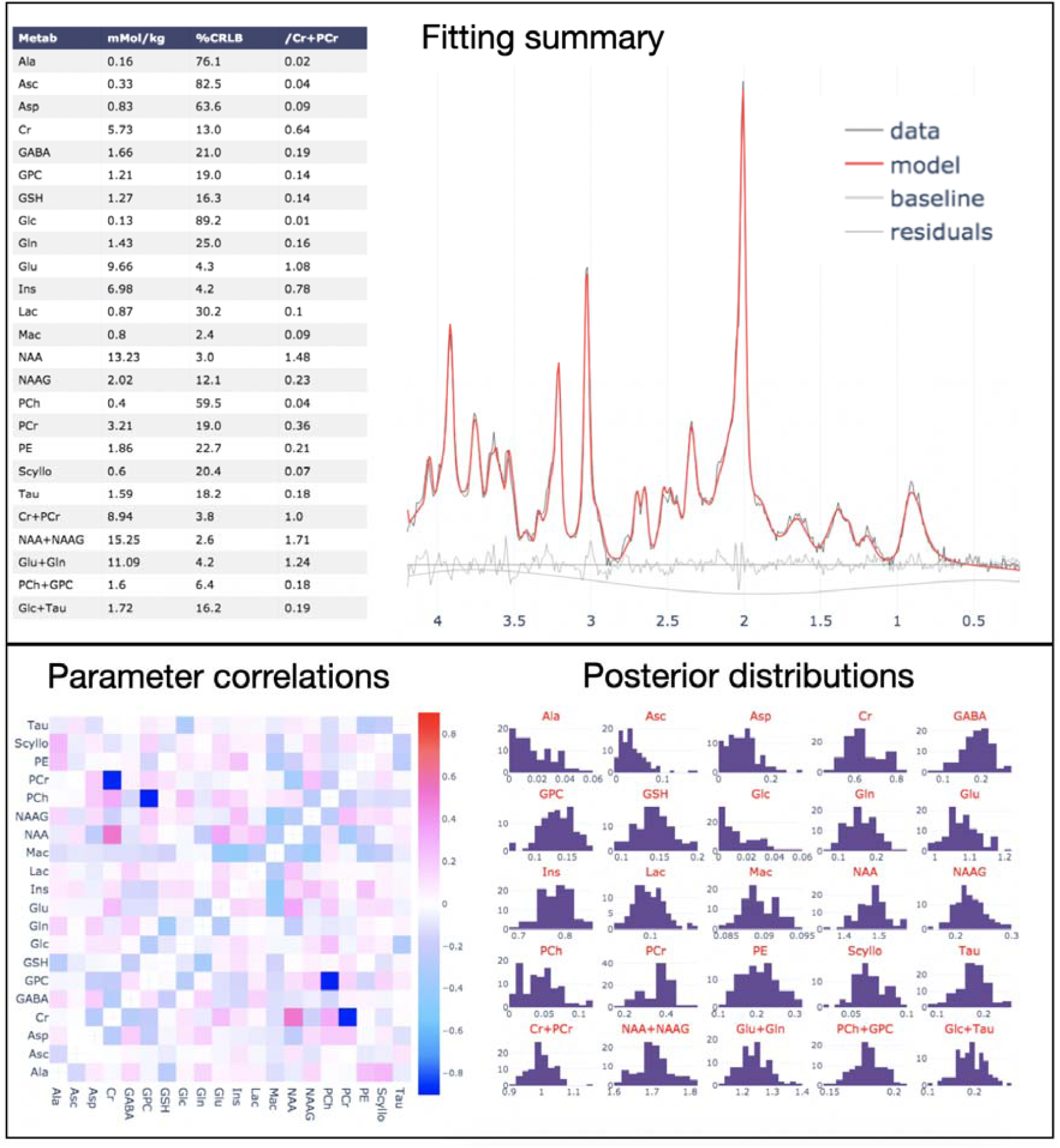
Extracts of the interactive HTML fitting report. **Top:** Metabolite concentrations summary and fit overlaid on data. Individual plots can be toggled on and off interactively. Bottom: correlations between metabolite concentrations from the Monte Carlo sampling (MCMC) and marginal posterior distributions of the metabolite concentrations. A full interactive fitting and pre-processing report is attached as supplementary information.

Visualisation of both time and frequency domain MRSI data alongside structural imaging data can be achieved using the FSL package tool FSLeyes (35) (see Figure 3).

**Figure 3 –.**
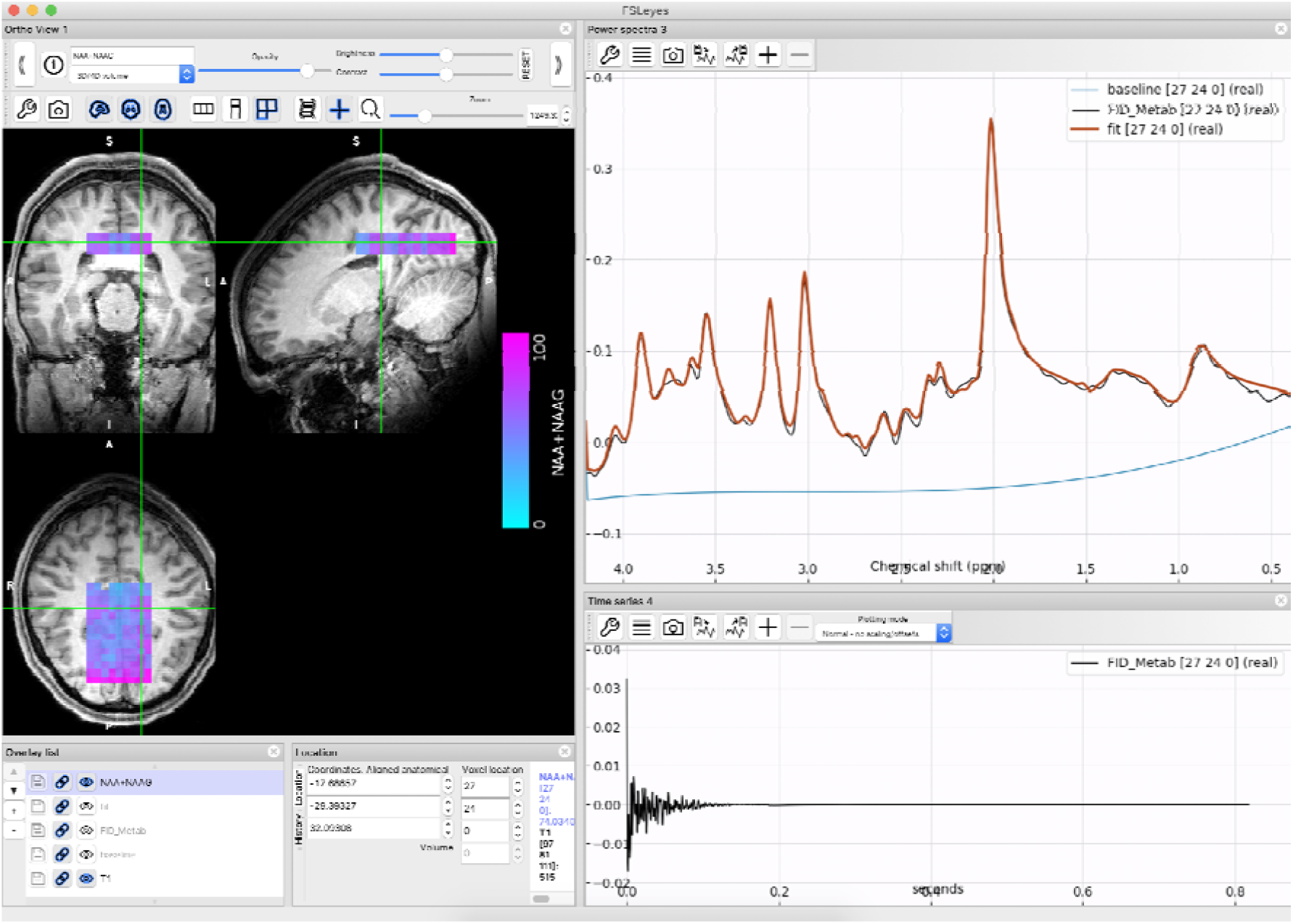
MRSI in FSLeyes. The results of processing and fitting of MRSI data are stored in 4D NIFTI files and can be viewed in a suitable viewer such as FSLeyes. Here a map of total NAA (NAA+NAAG) as measured using CONCEPT (Dataset 3) is overlaid on a T1 weighted image. In the right-hand side panel the real part of the time series data for the selected voxel is seen on the bottom, and on the top the real part of the spectral data is overlaid with the FSL-MRS fit and baseline estimate.

### Interactive FSL-MRS

In addition to the command line tools described in the previous sub-section, FSL-MRS may be run in an “interactive” way by loading the underlying python libraries into an interactive IPython environment. The same functionality and reporting interfaces that are available on the command-line are also available interactively. In this way FSL-MRS allows prototyping of new processing pipelines and tools whilst also providing familiarity for users of interactive scripting languages.

### Bayesian fitting

FSL-MRS implements linear combination modelling for fitting of basis spectra to data using Bayesian statistics to find an optimal solution. This method of fitting is robust whilst also outputting full posterior distributions of fitted metabolite concentrations to estimate concentration covariances and uncertainties.

The fitting module contains a standard fitting model appropriate for the fitting of a single independent spectrum. However, the fitting framework can accept an arbitrary forward model.

### Standard fitting model

The model for the complex-domain spectrum is

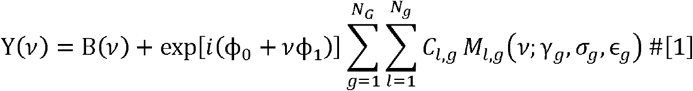

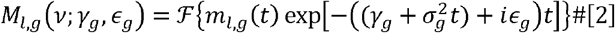

Where *v* denotes frequency, B(*v*) describes an n^th^-order complex polynomial estimate of the baseline, the second term applies a global 0^th^ and 1^st^ order phase and the final term is the sum of all scaled, shifted and broadened metabolite basis spectra *M*_*l,g*_(*v*; *γ*_*g*_, *σ*_*g*_, *ϵ*_*g*_). To avoid overfitting there is no flexibility in the metabolite line-shapes beyond shifting (ε) and broadening (γ, σ), which can be flexibly applied to *N*_G_ groups of metabolites (where each metabolite belongs to one and one only group). *ℱ* is the Fourier transform and m_l,g_(*t*) is the inverse Fourier transform of *M*_*l,g*_(*v*; 0,0,0).

No prior information or constraints on relative metabolite scaling is incorporated. The default polynomial baseline is second-order, but can be specified (or disabled entirely) by the user.

### Optimisation

Initialization is achieved using the truncated Newton algorithm as implemented in the SciPy package (36,37). The final fit is carried out over all model parameters using Metropolis-Hastings (a Markov Chain Monte Carlo algorithm) (18). The truncated Newton initialization can be used independently of the subsequent MCMC fit to provide a fast point-estimate of the metabolite concentration. In this work and in the summary reports generated by FSL-MRS, point-estimates of the metabolite concentrations from the MCMC algorithm are the arithmetic mean of the posterior distribution.

The forward model in Eq 1. is combined with an additive Gaussian white noise to produce the Likelihood function (which combines both real and imaginary part of the model prediction and data). The noise variance parameter is integrated out with a Jeffrey’s (1/x) prior. Priors on the concentration parameters are set to broad zero-mean half-Gaussians (i.e. with positivity constraint). Each of the line broadening parameters (γ, σ) are set to broad Gaussians (SD of 2.5 Hz) with a small positive centre (5Hz) and positivity constraints. Thus the prior is centred at an additional 10 Hz of line broadening in addition to the linewidth of the basis spectra. Shift and phase priors are set to broad Gaussians centred at zero with no additional constraints. Priors can be disabled (set to uniform) by the user. The baseline parameters are estimated in the initial nonlinear fitting then kept fixed in the MCMC stage. More details of the optimization choices (including initialisation, priors, and likelihood model) can be found in the Optimisation details section of the supporting information. Note that these details pertain to the results shown in this paper. The online software documentation will be kept up-to-date should these optimisation decisions change in future releases.

### Treatment of Macromolecular Signals

Macromolecular signal is observed as broad resonances in short echo-time spectra. The signal arises from amino acid residues of peptides. The methods described in “*Basis spectra simulation*” are not suitable for the creation of macromolecular basis spectra: macromolecular resonances are broad distributions of chemical shifts arising from many different peptide molecules rather than a single metabolite molecule. FSL-MRS therefore uses empirically measured macromolecular signal (e.g. from metabolite T1-nulled acquisitions) as a basis. The complex polynomial based baseline model is not designed to describe macromolecular signals.

For situations where empirically measured macromolecular signals are not available simulated basis spectra, generated at known chemical shift positions may be added to the set of basis spectra automatically. The details of these basis spectra at time of writing are listed in the supporting information (Supporting Information Table 5) and in the online documentation. Users may add additional peaks or modify the defaults. In all cases macromolecular basis spectra are treated identically to metabolite basis spectra, but are grouped separately to allow suitable separate optimisation of frequency shift and line broadening parameters.

### Validation of fitting

All methods in this work refer to version 1.0.5 of FSL-MRS.

### Simulation

Independently created simulated data was used to validate FSL-MRS. The simulated data was created by Malgorzata Marjanska, Dinesh Deelchand, and Roland Kreis for the ISMRM MRS Study Group’s Fitting Challenge (38). The data comprises 21 datasets (without artefacts) with varying SNR, linewidths, line shapes, metabolite concentrations and macromolecule content. Briefly: datasets 0-2 have increasing widths of Lorentzian line-shapes; 3-5 have increasing widths of Gaussian line-shapes; 6-9 vary the concentration of GABA/GSH; 10 has no macromolecular content; 11-13, 14-16; 17-19, and 20 have different spectral SNR (20, 30, 40 & 160 respectively). The data simulates a 3T PRESS sequence with a T_E_ of 30 ms.

Both water-suppressed and unsuppressed data is provided in an already pre-processed state. Basis spectra for 17 metabolites, including a macromolecular baseline, were provided by the challenge authors. The metabolites included are: alanine (Ala), ascorbate (Asc), aspartate (Asp), γ-aminobutyric acid (GABA), glucose (Glc), glutamine (Gln), glutamate (Glu), glutathione (GSH), glycine (Gly), myo-inositol (Ins), lactate (Lac), N-acetyl aspartate (NAA), N-acetyl aspartate glutamate (NAAG), phosphorylethanolamine (PE), scyllo-inositol (sIns), taurine (tau). In the analysis the following metabolites were treated together: NAA+NAAG, Glu+Gln, GPC+PCho, Cr+PCr, Glc+Tau, Ins+Gly. True concentration values for each metabolite in each dataset were supplied by the Fitting Challenge authors in a private communication.

Fitting was assessed for both the Newton and MCMC algorithms. The polynomial baseline was restricted to 0^th^ order and fitting was carried out between 0.2 and 4.2 ppm. After fitting, scaling of the raw metabolite concentrations was carried out using the unsuppressed water data, and concentration values were scaled accounting for provided tissue volume fractions (32).

Fitting performance was assessed using the mean and median percentage difference and absolute concentration difference from the true concentration values for each metabolite in all datasets. In addition, a summary statistic for each metabolite in each spectrum was calculated from the MCMC estimated posterior distribution as follows:

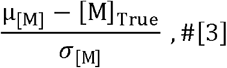

where *μ*_[*M*]_, *σ*_[*M*]_ are the mean and standard deviation of the fitted concentration of metabolite M and [M]_True_ is its true value. Intuitively this statistic can be interpreted as “how many standard deviations away from the true value is our estimate”.

### Phantom

Validation of unsuppressed, water scaled concentrations was carried out in a uniform aqueous phantom (SPECTRE, Gold Standard Phantoms, London, UK) containing six metabolites (NAA, Cr, Cho, Ins, Glu and Lac) using a previously published STEAM sequence at 7T (39,40). The sequence parameters were: 11 ms TE, 32 ms mixing time (T_M_), 10 s repetition time, 4096 samples, 6000 Hz bandwidth. Basis spectra were created using FSL-MRS. Basis spectra simulation used fully described (non-ideal) pulse shapes, gradients and timing parameters, and were conducted using a spatial resolution of 30 points in each gradient dimension. The concentrations of six metabolites was determined from 5 Hz exponentially line-broadened spectra from the phantom. This broadening was introduced to permit the use of the standard in vivo Bayesian priors in the optimisation. An additional doublet near 1.4 ppm was observed in the spectrum. It was established to be contaminant of the lactate feedstock used to create the phantom. It was fitted as alanine and included in the lactate concentration. Absolute concentrations were calculated from referencing the integral of the scaled creatine spectrum to an unsuppressed water spectrum taken to be equivalent to 55.5 M H_2_O. T_2_s were estimated from water and an average of metabolite singlet linewidths and concentrations were scaled for metabolite and water T_2_ relaxation.

### In vivo

FSL-MRS fitting was validated against LCModel (Version 6.3-1M) (13) in three in vivo datasets. The datasets covered different brain regions, sequences, and field strengths and are summarised in Table 2. Datasets one and two are single voxel sequences using STEAM and SPECIAL (41) sequences respectively and dataset three is 2D multi-voxel MRSI data collected using density-weighted CONCEPT with a semi-LASER volume selection module (2). STEAM and SPECIAL data were processed using *fsl_mrs_proc* respectively.

All subjects in these datasets were recruited in a manner approved by the appropriate Research Ethics Committee for each originating study (see references in Table 2).

Identical basis spectra were used in both FSL-MRS and LCModel. Basis spectra for datasets one and two were created in FSL-MRS using fully described RF pulses and gradients, coherence filtering and were simulated with 30 spatial points in each gradient dimension. The basis spectra comprised 19 and 17 simulated metabolites respectively to match previous analyses. Previously measured macromolecular spectra from metabolite-inversion-nulled sequences were included in the basis spectra. For dataset three existing basis spectra (as described in Ref (43)) were used. They were simulated in the simulation module of VeSPA (Versatile Simulation, Pulses and Analysis)(44) and comprise 19 simulated metabolites. Macromolecular spectra were not included; instead eight LCModel or FSL-MRS-simulated Gaussian macromolecule resonances were included in the analysis at the following positions: 0.91, 1.21, 1.43, 1.67, 1.95, 2.08, 2.25 and 3.00 ppm. For all datasets default “Concentration Ratio Priors” (also referred to as soft constraints) for metabolites were specified for the LCModel fit. I.e. the LCModel control file parameter “NRATIO” was set to the default value of 12 (corresponding to the first 12 ratio priors specified in §11.8 of the LCModel manual, of which numbers 8-12 are active in datasets 1&2 and numbers 1-6 & 8-12 are active in dataset 3). In LCModel the baseline flexibility parameter DKNTMN was set to 0.25, slightly above the default (0.15), whilst in FSL-MRS the baseline order was set to 2^nd^-order, 2^nd^-order and 4^th^-order for datasets one, two and three respectively.

Datasets one and two were fitted using LCModel and FSL-MRS (MCMC algorithm). Dataset three was fitted in LCModel and FSL-MRS (Newton algorithm) for speed. Highly correlated peaks (correlation coefficient⍰< ⍰−0.5) were combined (e.g. Cr+PCr, NAA+NAAG, PCh+GPC, Glu+Gln). Metabolite concentrations were expressed as a ratio to total creatine (Cr+PCr) and as molality concentrations using unsuppressed water as an internal reference. T_2_ relaxation was accounted for, but the unsuppressed water peak was assumed to correspond to pure water as anatomical images for tissue segmentation were not available for all datasets.

Data was compared voxel-wise for each metabolite in each dataset using the Pearson correlation coefficient and Bland-Altman bias and limits of agreement (45). Bias and limits of agreement were calculated in concentration units or as ratios to total creatine (Cr+PCr). They were summarised by expressing the bias as a percentage of the mean concentration value, and the limits of agreement as the width of the 95% confidence intervals expressed as a percentage of the mean value before averaging these values across all metabolites. In the comparisons, data was excluded if the estimated percentage Cramer Rao lower bounds on the metabolite concentrations exceeded 100% for either FSL-MRS or LCModel, or if the fitted value was more than four standard deviations from the mean value for that metabolite in that dataset. Metrics were calculated for both the water-scaled concentrations (water) and metabolite ratios (tCr); for all metabolites excluding the combined values (all) and for all metabolites including combined values but excluding those which were combined (combined).

## Results

### Output and reports

Figure 2 shows extracts of an example FSL-MRS fitting report. The extracts include a summary of the fit and metabolite concentrations, MCMC-estimated correlations between metabolite concentrations, and visualisations of the MCMC estimated distributions of the metabolite concentrations. Example fully interactive HTML reports for both fitting and processing are attached as supporting information. The same reports can be generated from example data included in the FSL-MRS package.

Figure 3 shows the results of fitting an MRSI grid of voxels from a single density weighted CONCEPT from dataset three (Table 2). NIfTI format viewers such as FSLeyes can be used to simultaneously view anatomical images, fitted metabolite concentrations, the spectral data and the FSL-MRS fit. In Figure 3 the total NAA concentrations are overlaid on a T1w image centred around the calcarine sulcus.

Fitting results may be exported in NIfTI or comma separated value (CSV) format or carried forward in python for further analysis.

### Validation

#### Simulation

Figure 4 summarises the results of the validation on simulated data for all metabolites in all simulated datasets. Detailed plots for each metabolite are included as Supporting Information Figure S1.

**Figure 4 –.**
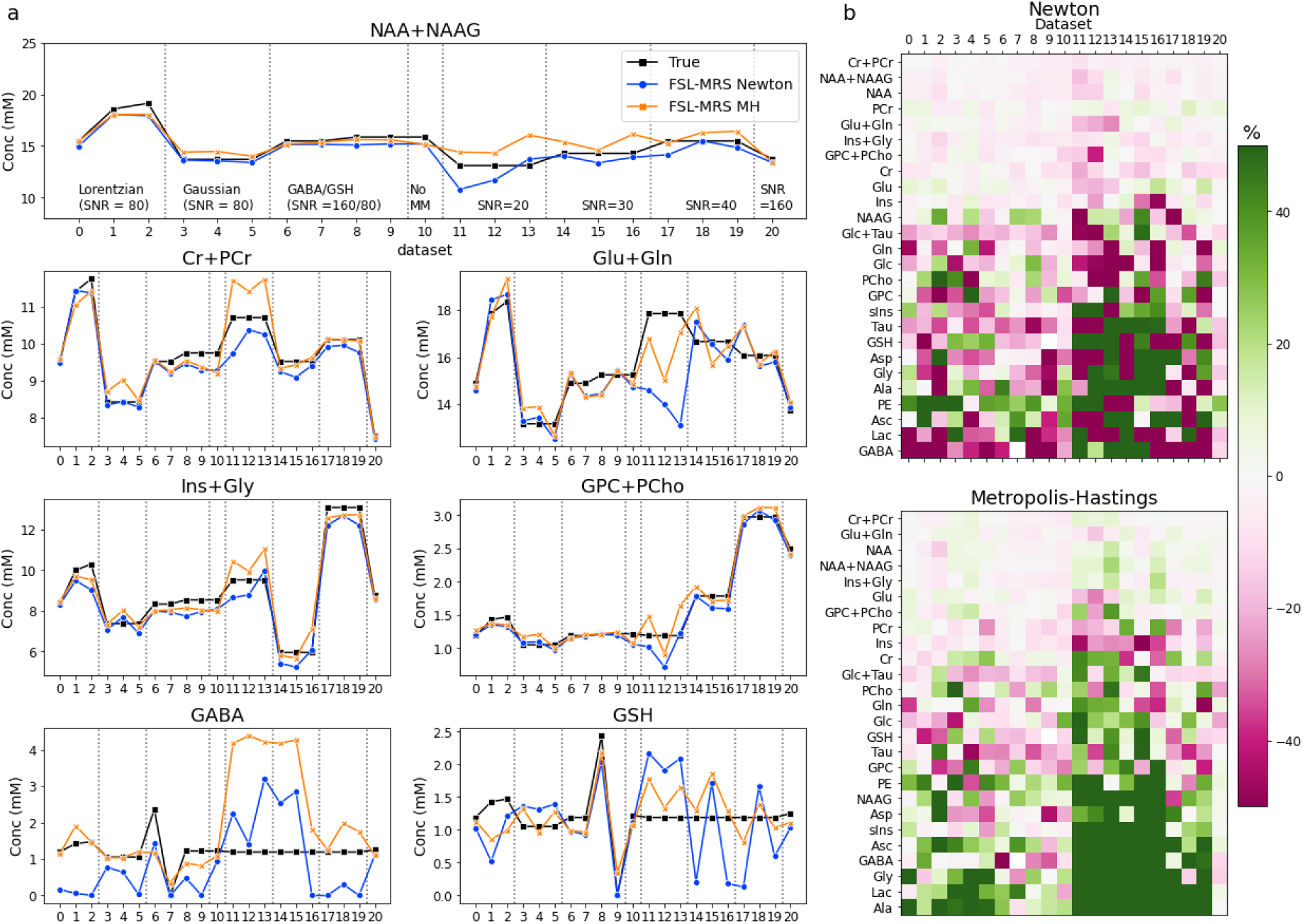
Simulation validation. **a** Comparison of FSL-MRS measured concentrations for each MRS fitting challenge dataset for seven metabolites. Equivalent plots for all metabolites can be found in Supporting Information Figure S1. **b** Percentage difference from true values for all metabolites for all datasets. The metabolites are sorted by mean difference. Both fitting algorithms (Newton [top] & Metropolis Hastings [bottom]) are shown.

For all metabolites across all 21 datasets the Newton algorithm achieved a mean (median) absolute concentration difference of 0.60 (0.41) mM and a mean (median) absolute percentage difference of 30.6 (14.9) %. And for the MCMC algorithm 0.60 (0.37) mM and 35.2 (11.9) %. For the five most prominent signals (NAA+NAAG, Cr+PCr, Glu+Gln, Ins+Gly, GPC+PCho) the MCMC algorithm had a mean difference of 0.48 mM or 5.4%. The mean (±SD) number of standard deviations from the true value (Equation 3) was 0.57±0.43. 98.9% of true metabolite concentration values were between the 5^th^ and 95^th^ percentiles of the MCMC estimated posterior distributions. Uncombined choline (GPC & PCh) and creatine (PCr&Cr) peaks were excluded from the calculation as described in the original fitting challenge results.

#### Phantom

Figure 5 summarises the results of the absolute concentration validation in phantom. The mean absolute percentage difference from the true concentration across all metabolites was 3.39% (range −7.1% [Lac] to 1.1% [Glu]).

**Figure 5 –.**
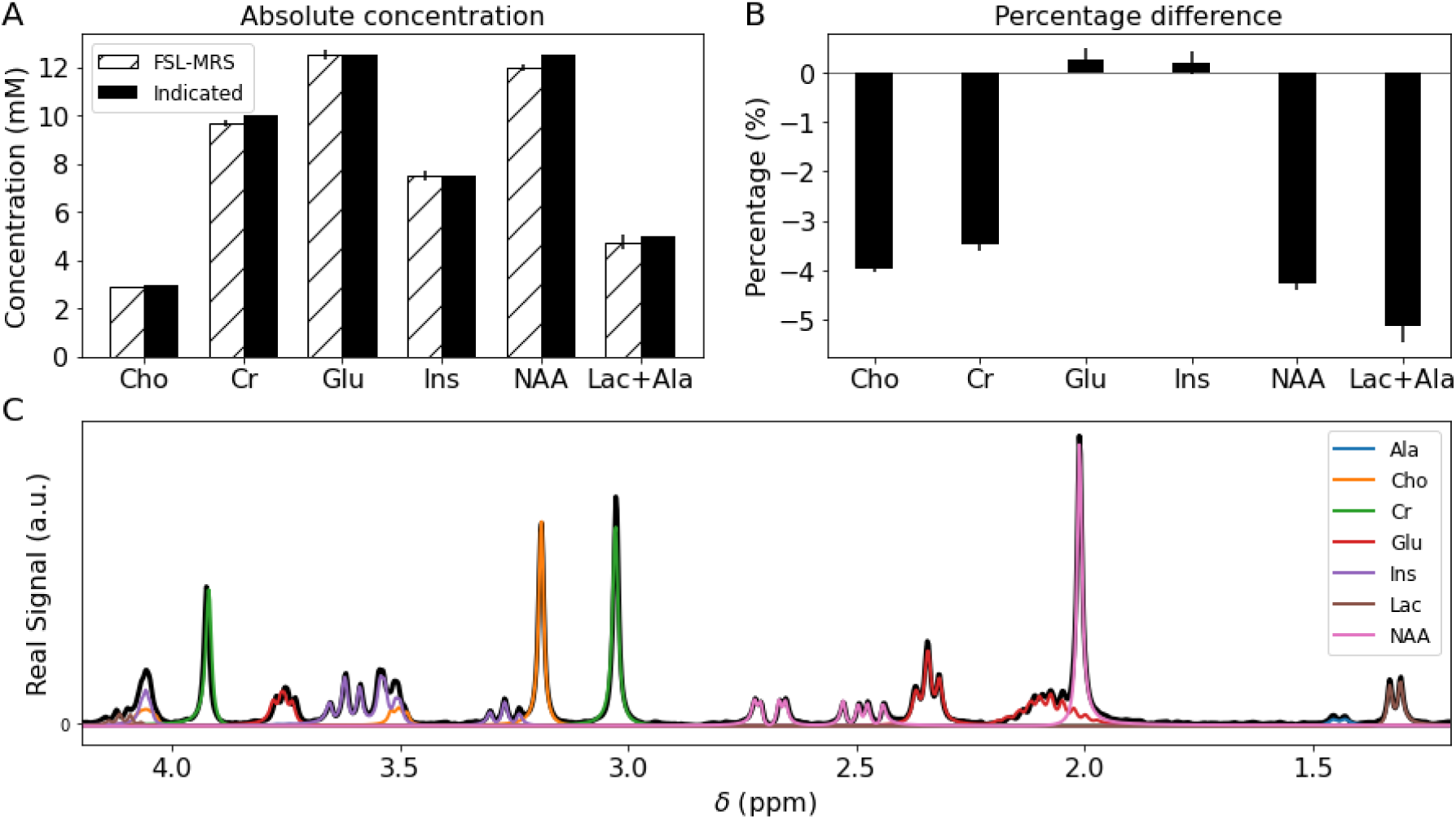
Phantom validation. A) Absolute concentration of fitted metabolite compared to known concentrations. CRLB are indicated by vertical bars. B) Percentage difference from true value. C) Data overlaid with FSL-MRS fit, each metabolite fit is shown in a different colour. The doublet at 1.4 ppm is fitted as ‘Ala’ and included with lactate.

The single creatine basis set used was unable to simultaneously fit both creatine singlet peaks (CH2 at 3.93 ppm, CH3 at 3.03 ppm) with small residuals. The creatine singlets have been observed to have different T2 relaxation properties which is unmodeled in these basis spectra (46), and could account for the observed difference in fit quality between peaks.

#### In vivo

Table 3 summarises the in vivo fitting validation correlations and Bland-Altman metrics for each dataset. The per-metabolite correlations for each of the three datasets and for both referencing method is provided in the Supporting Information (Table 6).

Mean correlations between FSL-MRS and LCModel in all data sets achieved a correlation over 0.5, and correlations were similar for both water-scaled concentrations and metabolite ratios. Correlations for the combined metabolite group were higher than the uncombined ‘all’ group, as the high SNR combined metabolites (Cr+PCr, PCh+GPC, Glu+Gln, NAA+NAAG, Ins) achieved correlations in the range 0.81 – 0.98 for all datasets. The highest metabolite correlation across all three datasets was achieved by total choline (0.85), the lowest was glucose (0.34), the median per-metabolite correlation was 0.70. Figure 6 shows scatter plots for a sample of metabolites for each dataset (for display purposes only, concentrations and ratios were normalised to the maximum value fitted by either LCModel or FSL-MRS). The scatter plots for all metabolites for each dataset are included as supplementary information (Supporting Information Figures S2-4).

**Figure 6 –.**
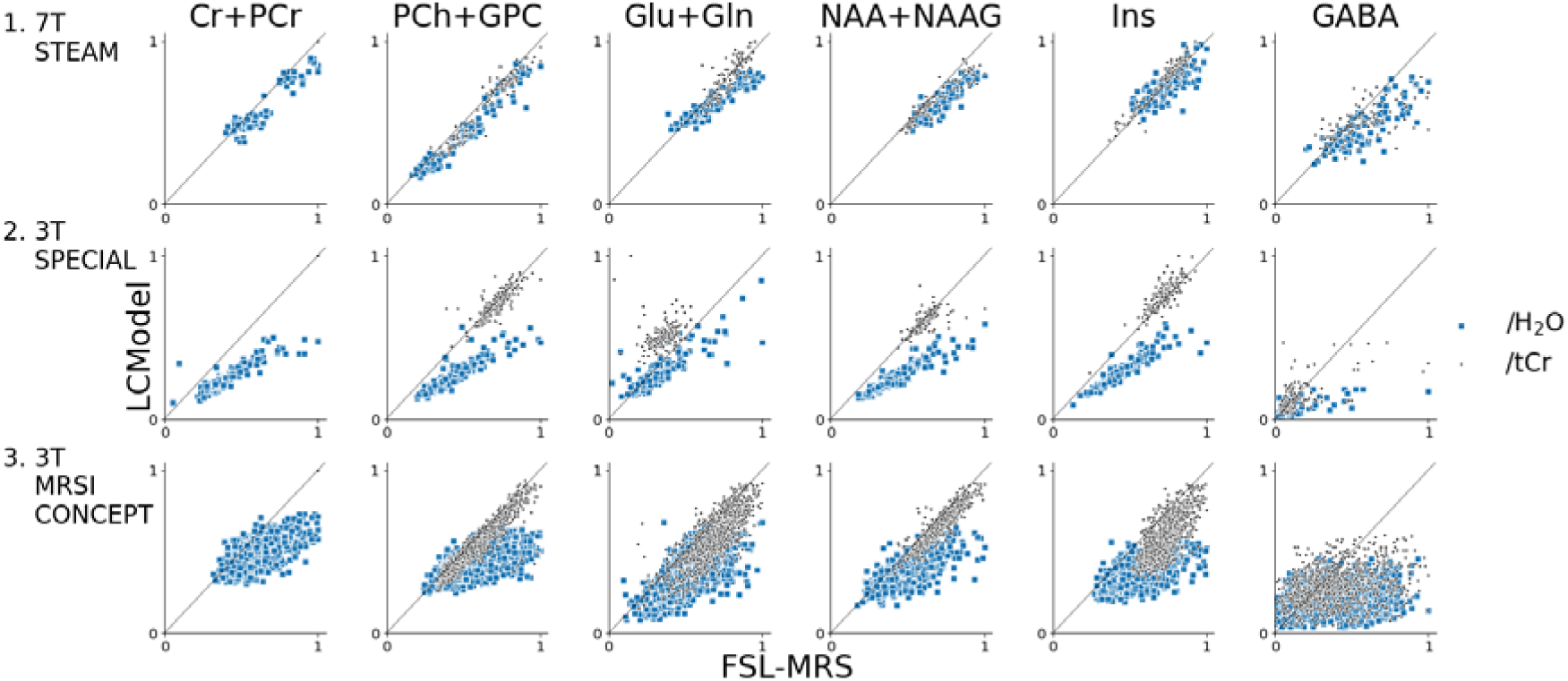
Summary of *in vivo* validation. Correlation plots of a selected group of metabolites for each validation dataset. Solely for display purposes, ratios to unsuppressed water and total creatine (Cr+PCr) are normalised to the maximum value fitted by either FSL-MRS or LCModel. Correlation plots for all metabolites are shown in the supporting documents.

When averaged across all metabolites Bland-Altman metrics showed a consistent bias for higher metabolite concentrations (mean of 14% to 37%) and metabolite ratios (11% to 24%) in FSL-MRS compared to LCModel. Bias was higher for water-scaled concentrations compared to metabolite ratios and lower for the combined metabolites. Bland-Altman plots are shown for high SNR metabolites (Cr+PCr, PCh+GPC, Glu+Gln, NAA+NAAG, Ins) in Supporting Information Figure S5. These metabolites showed much lower bias when referenced to total creatine (Supporting Information Table 7) vs water referencing (Supporting Information Table 8).

## Discussion

FSL-MRS is an end-to-end spectroscopy analysis package. It is designed to be used flexibly: either implementing all stages of the MRS analysis pipeline or being used as a modular part of another pipeline. The package is scriptable on the command line requiring no interaction – suitable for analysis of large datasets and for deployment with high performance computing, or it can be used interactively – for pipeline prototyping and novel analyses.

It achieves modularity by operating on data stored in a standard file type, NIfTI, which is already in use throughout neuroimaging. Existing packages for handling NIfTI data exist in many programming languages (e.g. NiBabel in Python and the “image processing toolbox” in Matlab) enabling FSL-MRS to be integrated with other MRS analysis programs. Results generated in NIfTI format allows straightforward integration of MRS data into multi-contrast analysis in existing neuroimaging toolboxes (e.g. FSL). Both FSL-MRS and Python are open source and free for academic and non-commercial use.

The package includes visualisation modules for generating interactive HTML reports viewable in a wide range of internet browsers. Visualisation of data and fitting results can also be accomplished in NIfTI format viewers due to the use of standard data types. Visualisation of data remains important whilst fully automatic quality control of MRS data remains not widespread (20).

Validation of the FSL-MRS fitting module was carried out on simulation, phantom and in vivo data. Validation on the simulated data showed low absolute concentration errors except in those datasets with low spectral SNRs (20 & 30) and in peaks with low SNR and high correlation with neighbours (e.g. GABA). MCMC fitting of metabolites with low concentrations generate skewed distributions which are not well described with a single point-statistic (the mean value) which may contribute to the marginally better performance of the Newton algorithm. Phantom validation indicated that the package correctly implements calculation of absolute concentrations using scaling to unsuppressed water in the case of pure water.

In in vivo data the validation was against LCModel, an established and widely used fitting program. Bias towards higher metabolite ratios in FSL-MRS was observed for water-scaled concentrations and, to a lesser extent, for relative metabolite ratios. The latter might arise from FSL-MRS not implementing priors between relative metabolite concentrations, a default setting in LCModel which was enabled in this analysis. Soft constraints in LCModel restrict certain metabolite concentration ratios (ratios of low SNR metabolites to a weighted average of NAA, total creatine and total choline) to be within a certain normally distributed range. The larger differences in water-scaled metabolite concentrations is likely due to the different implementations of flexible baselines in the two packages. LCModel and other programs (47,48) implement a spline-based baseline, in this work we have chosen a complex polynomial implementation. The inclusion of a 0^th^-order term allows a uniform vertical shift in baseline across all frequencies, not typically possible with spline-baselines.

Across all voxels in the MRSI dataset negative correlations were observed between absolute concentrations and the 0^th^ order polynomial baseline parameters. A baseline below zero will increase the reference peak’s absolute integral and result in a large ratio when compared with the integral of unsuppressed water. The effect of the precise implementation of flexible baselines on metabolite concentrations in fitting packages is complex (49,50) with dependence on acquisition, description of macromolecules in basis spectra and optimisation algorithm. FSL-MRS’s implementation of a complex polynomial baseline does not offer a solution to this complexity, but the implementation is simple to understand and implement, is unlikely to cause overfitting, and is only parametrised by 2n nuisance parameters for an order-n baseline. If enabled, the MCMC algorithm enables the user to calculate the covariance of the baseline parameters with the metabolite concentrations. An example MCMC correlation matrix of a single spectrum from dataset #1 (7T STEAM) including baseline parameters shows baseline parameters only correlate strongly with the macromolecule concentration (Supporting Information Figure S6). Efforts to widely measure and account for differences in fitting software (51) will be essential to provide program quality assurance and allow for meaningful use of pooled data analysed using different tools.

Fitting using the MCMC algorithm allows the user to generate the full posterior distribution for each fitted parameter, including metabolite concentrations. This information is essential to understanding the uncertainties inherent in the estimation of the parameters. It also offers the opportunity to carry forward this information into subsequent study analysis, reducing the need for arbitrary quality cut-offs to be used. However, fitting using the MCMC algorithm is inherently slower than methods that only provide point estimates, taking 10s of seconds rather than seconds to compute the results for each voxel. It may be possible to achieve the estimation of the posterior distributions in the time frame of a few seconds using a variational inference optimiser, which is under development (52).

Operation of the package still requires the user to provide expert knowledge in two places: data conversion and generation of basis spectra. At the data conversion stage, the user must either use a file format understood by *spec2nii* and must interpret the structure of the data within that format, or provide a full conversion, including orientation information, for their own data format. Generating correct basis spectra requires the user to provide an accurate description of the RF pulses, timings and gradients in the localisation module of their sequence. Documentation for the package has been created to mitigate difficulties in these stages. *fsl_mrs* (SVS fitting) can interpret a select few other formats (LCModel “.RAW” & jMRUI “.txt”).

The FSL-MRS MCMC fitting module accepts an arbitrary forward model. In future work we intend to use this framework to investigate the advantages of fitting multiple spectra simultaneously with a specialist model (e.g. for diffusion-weighted, edited or functional MRS). The FSL-MRS package is under continued development and refinement, online documentation provides the latest and up-to-date information on the package. Currently the package is optimised for 3T and 7T in vivo human 1H-MRS data, fitting routines, basis spectra and prior knowledge will need to be suitably modified for a greater range of data.

## Conclusion

We have presented a new end-to-end spectroscopy processing package incorporating Bayesian fitting of spectra. The package is open-source, modular and freely available. In this work validation of the package by simulation, in phantom and in three in vivo datasets has been provided. The complete package is available for download at git.fmrib.ox.ac.uk/fsl/fsl_mrs, through the open source package management system Conda (Continuum Analytics, Inc), and will be available as part of FSL (fsl.fmrib.ox.ac.uk).

## Supporting information

Interactive SVS fitting report

Interactive SVS processing report

## Acknowledgements

Saad Jbabdi is funded by a Wellcome Trust Collaborative Award (215573/Z/19/Z).

Charlotte Stagg is supported by the Wellcome Trust and the Royal Society (102584/Z/13/Z).

The Wellcome Centre for Integrative Neuroimaging is supported by core funding from the Wellcome Trust (203139/Z/16/Z).

We thank Phil Cowen and Beata Godlewska for providing the STEAM data, collection of which was funded by the MRC [MR/K022202/1].

We thank Sana Suri, Enikő Zsoldos, and Klaus P. Ebmeier for providing data from the Whitehall II MRI substudy funded by the UK Medical Research Council (MRC) grant [G1001354; ClinicalTrials.gov Identifier: NCT03335696]. The research was also supported by the EU Horizon 2020 [Grant agreement number: 732592; “Lifebrain”] and the HDH Wills 1965 Charitable Trust [1117747].

We thank Aislin Sheldon, Holly Bridge and Jasleen Jolly for providing the MRSI validation data. This data was collected using funding from the MRC [MR/K014382/1], National Institute for Health Research (NIHR) [CA-CDRF-2016-02-002] and an MRC studentship.

We thank Malgorzata Marjanska, Dinesh Deelchand, and Roland Kreis for generating and providing the MRS Fitting challenge data.

We thank Sean Fitzgibbon, Michiel Cottaar and Paul McCarthy for providing their assistance and expertise in all matters Python.

We thank Jamie Near and Uzay Emir for useful discussions throughout the development of FSL-MRS.

## Figure captions

Figure 1 – FSL-MRS organisation and workflow. Raw data in proprietary or other formats are converted to NIfTI by spec2nii. Processing can then be carried out in stages, operating on NIfTI files, using fsl_mrs_proc, or in a single python script fsl_mrs_preproc for standard SVS sequences. Basis spectra can be generated for fitting using fsl_mrs_sim given a json description for the sequence. Fitting and quantitation are then carried out by fsl_mrs and fsl_mrsi as appropriate. Interactive HTML reports are generated for viewing in the user’s browser. Spectroscopy data in NIfTI format can be viewed overlaid with other MR contrasts in FSLeyes.

Figure 2 – Extracts of the interactive HTML fitting report. **Top:** Metabolite concentrations summary and fit overlaid on data. Individual plots can be toggled on and off interactively. Bottom: correlations between metabolite concentrations from the Monte Carlo sampling (MCMC) and marginal posterior distributions of the metabolite concentrations. A full interactive fitting and pre-processing report is attached as supplementary information.

Figure 3 – MRSI in FSLeyes. The results of processing and fitting of MRSI data are stored in 4D NIFTI files and can be viewed in a suitable viewer such as FSLeyes. Here a map of total NAA (NAA+NAAG) as measured using CONCEPT (Dataset 3) is overlaid on a T1 weighted image. In the lower panel the real part of the time series data for the selected voxel is seen on the left, and on the right the real part of the spectral data is overlaid with the FSL-MRS fit and baseline estimate.

Figure 4 – Simulation validation. **a** Comparison of FSL-MRS measured concentrations for each MRS fitting challenge dataset for seven metabolites. **b** Percentage difference from true values for all metabolites for all datasets. The metabolites are sorted by mean difference. Both fitting algorithms (Newton [top] & Metropolis Hastings [bottom]) are shown.

Figure 5 – Phantom validation. A) Absolute concentration of fitted metabolite compared to known concentrations. CRLB are indicated by vertical bars. B) Percentage difference from true value. C) Data overlaid with FSL-MRS fit, each metabolite fit is shown in a different colour. The doublet at 1.4 ppm is fitted as ‘Ala’ and included with lactate.

Figure 6 – Summary of *in vivo* validation. Correlation plots of a selected group of metabolites for each validation dataset. Solely for display purposes, ratios to unsuppressed water and total creatine (Cr+PCr) are normalised to the maximum value fitted by either FSL-MRS or LCModel. Correlation plots for all metabolites are shown in the supporting documents.

## Supporting Information

In FSL_MRS_Supporting_Info.docx:

1. A description of Optimisation details including MCMC fitting.
2. Supporting Information Table 1. File formats supported by spec2nii.
3. Supporting Information Table 2. Quantification constants: tissue water density.
4. Supporting Information Table 3. Quantification constants: T1 values.
5. Supporting Information Table 4. Quantification constants: T2 values.
6. Supporting Information Table 5. Synthetic macromolecular basis spectra specification.
7. Supporting Information Table 6. In vivo validation per-metabolite Pearson correlations.
8. Supporting Information Table 7. Bland-Altman statistics for creatine-referenced high-SNR metabolite peaks.
9. Supporting Information Table 8. Bland-Altman statistics for water-referenced high-SNR metabolite peaks.
10. Supporting Information Figure S1: Simulation validation results for all metabolites.
11. Supporting Information Figure S2: All metabolite correlations for dataset 1 (STEAM, 7T).
12. Supporting Information Figure S3: All metabolite correlations for dataset 2 (SPECIAL, 3T).
13. Supporting Information Figure S4: All metabolite correlations for dataset 3 (MRSI, 3T).
14. Supporting Information Figure S5: Bland-Altman plots for selected metabolites.
15. Supporting Information Figure S6: MCMC parameter correlations of a single dataset. As separate files:
16. Fitting_report.pdf: Static example of an FSL-MRS fitting report for a single-voxel dataset. Interactive version available at users.fmrib.ox.ac.uk/∼saad/fsl_mrs/reports/fitting_report.html.
17. Processing_report.pdf: Static example FSL-MRS processing report for a single voxel dataset. The interactive version is available online at users.fmrib.ox.ac.uk/∼saad/fsl_mrs/reports/fitting_report.html.

