## Supplemental tables and figures for "FSL-MRS: An end-to-end spectroscopy analysis package"

Supporting information to “FSL-MRS: An end-to-end spectroscopy analysis package”

### Optimisation details:

Prior to fitting, both the data (FID) and the basis are normalised such that the norm of the FID and the norm of the mean basis spectrum equals 100 (by default). This improved robustness to the range of data that we have encountered where the scaling can arbitrarily span many orders of magnitude. If a water FID is included for absolute quantitation, the same scaling factor is applied to both metabolite FID and water FID. This scaling also means that we can use priors on the concentration parameters without worrying about the scaling of the input data.

All model parameters are optimized jointly in the Truncated Newton optimisation, including the baseline coefficients, the global phase parameters, the line shape (shifting and broadening) parameters, and the concentrations. This optimisation uses no prior constraints except for positivity of a subset of the parameters (line shape [gamma/sigma] and concentrations).

The algorithm does require initialisation, which is particularly tricky for the line shape parameters. We use the following strategy for initialisation:

+ The global phase parameters $\Phi_{0}$ and $\Phi_{1}$ are initialised to zero. Although this may not be an optimal starting point for real-life spectra, we note that the tool allows the user to apply a phase correction to the spectrum prior to the fitting. This is done by maximizing the power in the real part of the spectrum within the fitting range (default: 0.2-4.2 ppm).

+ The line shape parameters are initialised using a nonlinear optimisation procedure that assumes a single value for these parameters for all metabolites (i.e. there is a single metabolite group) and uses the Powell (gradient-free) method as implemented in Scipy. The cost function of this optimisation is coded in the following way (pseudo-code):

Def Loss($\epsilon$, $log(\gamma)$, $log(\sigma)$):

+ $\gamma=\exp\left( \log\gamma\right), \sigma=\exp(\log\sigma)$

+ shift and broaden basis spectra

+ concatenate basis spectra with baseline polynomials

+ calculate least-squares fit to the spectral data using Moore-Penrose pseudoinverse

+ set negative concentrations to zero

+ return sum squared error

The above loss function is minimized (with initial values (0, log(10^-5^), log(10^-5^))

Minimizing the above loss function gives initial values for epsilon, gamma and sigma. These are then used to initialise the concentrations and baseline parameters using least squares fitting and setting negative concentrations to zero.

#### MCMC fitting:

We assume the following forward model for the data:

$$Y=F\left( \Theta\right)+noise$$

where $\Theta$ is the set of all parameters (baseline, line shape, phase, concentration), and the noise term is zero-mean Gaussian $noise\sim N(0,\sigma_{n}^{2}I_{m})$ where $I_{m}$ is the identity matrix and $m$ the number of data points. This yields a likelihood function of the form:

$P\left( Y | \Theta,\sigma_{n} \right)=N(F\left( \Theta\right),\sigma_{n}^{2}I_{m})$.

We further assume a Jeffrey’s prior on the noise standard deviation $\sigma_{n}$:

$P\left( \sigma_{n} \right)=1/\sigma_{n}$.

This means that we can integrate out $\sigma_{n}$ (as it is a nuisance parameter) to obtain a likelihood function of the form:

$P\left( Y | \Theta\right)=\int P\left( Y | \Theta,\sigma_{n} \right)P\left( \sigma_{n} \right)d\sigma_{n}\propto{|Y-F\left( \Theta\right)|}^{-\frac{m}{2}}$,

where $\left| . \right|$ denotes the Euclidean norm.

By default, the algorithm uses 100 samples for burn-in (I.e. initialisation of the sampling) followed by 500 sampling iterations where 1 in 10 samples are retained to reduce between-sample correlations.

Finally, we also use the following Gaussian priors for $\Theta$ (in addition to positivity, i.e. truncated Gaussians, for $\gamma_{g},\sigma_{g}, C_{l,g}$:

$$P\left( \gamma_{g} \right)=N\left( 5\mathrm{Hz},2.5\mathrm{Hz} \right)$$

$$P\left( \sigma_{g} \right)=N(5\mathrm{Hz},2.5\mathrm{Hz})$$

$$P\left( \epsilon_{g} \right)=N\left( 0,0.05\mathrm{ppm} \right)$$

$$P\left( C_{l,g} \right)=N\left( 0,1 \right)$$

$$P\left( \Phi_{0} \right)=N(0,5\text{deg})$$

$$P\left( \Phi_{1} \right)=N\left( 0,10\mu s \right)$$

Optionally the user may adjust the currently defined priors by modifying the hard-coded values in the *fsl_mrs.utils.constants* module, or by setting a command line flag to make all priors uniform.

### Supporting tables and figures

| **Vendor** | **Format** | **File extension** | **SVS** | **MRSI** | **Automatic orientation** |
| --- | --- | --- | --- | --- | --- |
| Siemens | Twix | .dat | Yes | No | Yes |
| Siemens | DICOM | .dcm/.ima | Yes | Yes | Yes |
| Philips | SPAR/SDAT | .SPAR/.SDAT | Yes | No | Yes (no tests available) |
| GE | p-file | .7 | Yes | No | Yes (no tests available) |
| N/A | LCModel | .RAW/.H2O | Yes | No | No |
| N/A | jMRUI | .txt | Yes | No | No |
| N/A | ASCII | .txt | Yes | No | No |

Supporting Table 1. File formats supported by spec2nii.

| **Tissue** | **Tissue Water Density (g/ml)** |
| --- | --- |
| GM | 0.78 |
| WM | 0.65 |
| CSF | 0.97 |

Supporting Table 2. Quantification constants: tissue water density

| **Field strength (T)** | **T1 Water: WM (s)** | **T1 Water: GM (s)** | **T1 Water: CSF (s)** | **T1 Metabolites (s)** |
| --- | --- | --- | --- | --- |
| 3 | 0.97 | 1.50 | 4.47 | 1.29 |
| 7 | 1.21 | 2.05 | 4.43 | 1.43 |
| *References* | *1-6* | *1-6* | *4* | *2,7-9* |

Supporting Table 3. Quantification constants: T1 values. References are listed at the end of the supporting information.

| **Field strength (T)** | **T2 Water: WM (ms)** | **T2 Water: GM (ms)** | **T2 Water: CSF (ms)** | **T2 Metabolites (ms)** |
| --- | --- | --- | --- | --- |
| 3 | 73 | 88 | 2030 | 194 |
| 7 | 55 | 50 | 1050 | 131 |
| *References* | *1,3,10-11* | *1,3,10-11* | *12* | *7-9,13-15* |

Supporting Table 4. Quantification constants: T2 values. References are listed at the end of the supporting information.

| **Peak name** | **Peak location(s) (ppm)** | **Peak relative amplitude(s)** | **Peak broadening (gamma/sigma)** |
| --- | --- | --- | --- |
| MM09 | 0.9 | 3 | 10/20 |
| MM12 | 1.2 | 2 | 10/20 |
| MM14 | 1.4 | 2 | 10/20 |
| MM17 | 1.7 | 2 | 10/20 |
| MM21 | 2.08, 2.25, 1.95, 3.0 | 1.33, 0.22, 0.33, 0.4 | 10/20 |

Supporting Table 5. Synthetic macromolecular basis spectra specification. Values are correct for the current version (1.0.5) at time of publication. Gamma and sigma are given in internal calculation units.

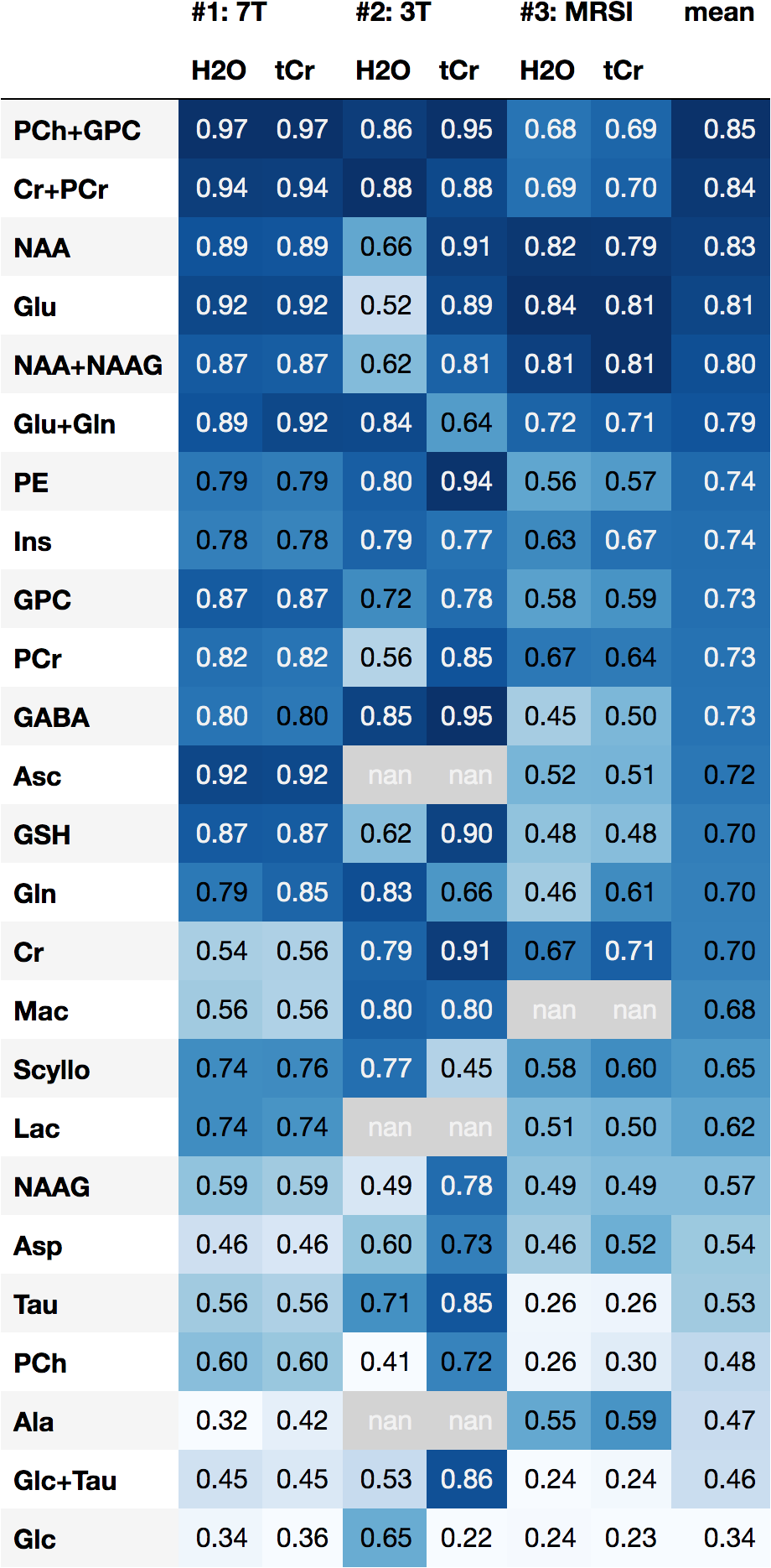

Supporting Table 6: Pearson correlation between LCModel and FSL-MRS for each metabolite. Highest correlations are shaded darkest, metabolites are listed highest average correlation to lowest. Metabolites absent from basis spectra sets are marked as “nan”.

|  | **7T** | | | **3T** | | | **MRSI** | | |
| --- | --- | --- | --- | --- | --- | --- | --- | --- | --- |
| **Metabolite** | *Mean value* | *Bias* | *LoA* | *Mean value* | *Bias* | *LoA* | *Mean value* | *Bias* | *LoA* |
| NAA+NAAG | 1.37 | 0.10 | 0.55 | 1.53 | 0.08 | 1.99 | 1.30 | 0.08 | 0.46 |
| Glu+Gln | 1.47 | -0.01 | 0.45 | 1.46 | -0.54 | 1.57 | 1.16 | 0.16 | 0.76 |
| PCh+GPC | 0.16 | 0.01 | 0.05 | 0.18 | 0.00 | 0.11 | 0.20 | 0.01 | 0.11 |
| Ins | 0.75 | -0.04 | 0.20 | 0.79 | -0.03 | 0.46 | 0.76 | 0.14 | 0.53 |
| Cr+Pcr | 1.00 | - | - | 1.00 | - | - | 1.00 | - | - |

Supporting Table 7. Bland-Altman statistics for creatine-referenced high-SNR metabolite peaks. Bias calculated as FSL-MRS minus LCModel. All values relative to tCr.

|  | **7T** | | | **3T** | | | **MRSI** | | |
| --- | --- | --- | --- | --- | --- | --- | --- | --- | --- |
| **Metabolite** | *Mean value* | *Bias* | *LoA* | *Mean value* | *Bias* | *LoA* | *Mean value* | *Bias* | *LoA* |
| NAA+NAAG | 14.53 | 2.04 | 4.94 | 13.78 | 6.35 | 12.03 | 12.96 | 3.43 | 11.11 |
| Glu+Gln | 15.87 | 1.37 | 7.67 | 13.48 | 1.98 | 13.13 | 11.76 | 3.77 | 12.75 |
| PCh+GPC | 1.85 | 0.26 | 0.82 | 1.61 | 0.83 | 1.46 | 1.95 | 0.53 | 1.94 |
| Ins | 7.88 | 0.23 | 3.57 | 6.99 | 3.00 | 5.20 | 7.51 | 2.96 | 7.30 |
| Cr+Pcr | 11.04 | 0.90 | 4.44 | 8.94 | 4.46 | 7.39 | 9.89 | 1.87 | 7.46 |

Supporting Table 8. Bland-Altman statistics for water-referenced high-SNR metabolite peaks. Bias calculated as FSL-MRS minus LCModel. All values in mM.

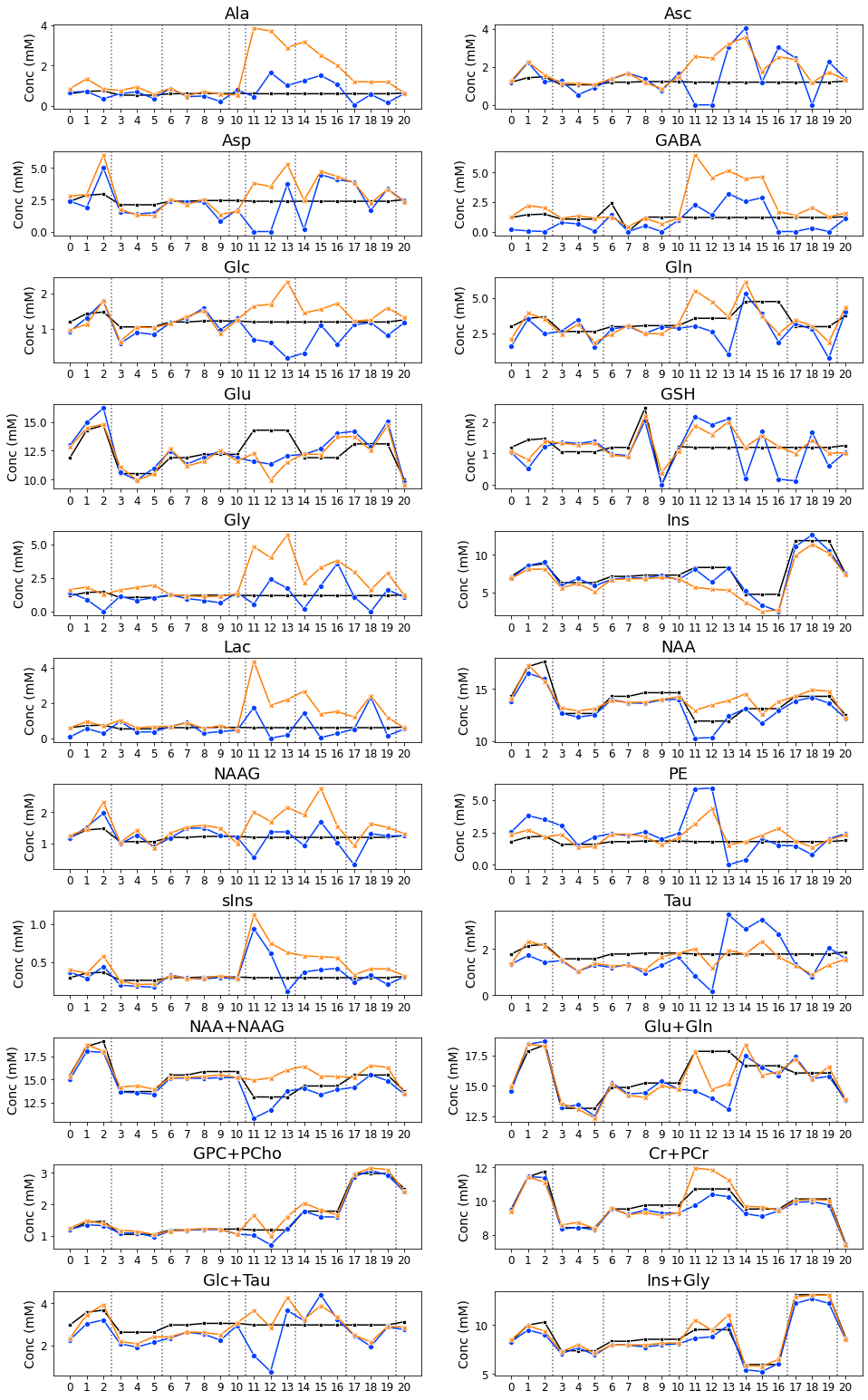

Supporting Figure 1: Simulation validation results for all metabolites, in the style of figure 4. True concentrations are marked as a black line, the Newton optimisation is in blue and the MH optimisation in orange.

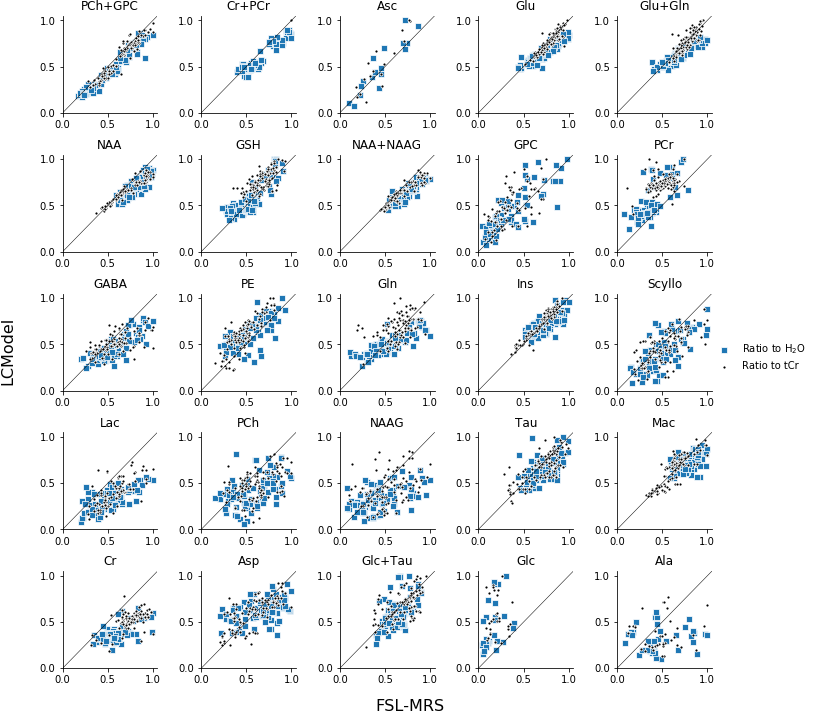

Supporting Figure 2: All metabolite correlations for dataset 1 (STEAM, 7T). Solely for display purposes, ratios to unsuppressed water and total creatine (Cr+PCr) are normalised to the maximum value fitted by either FSL-MRS or LCModel.

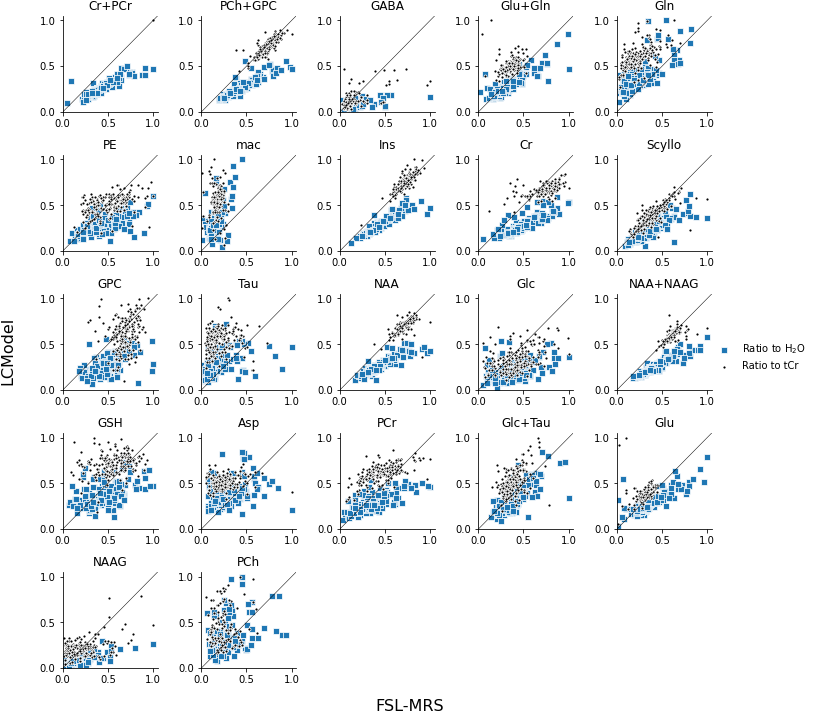

Supporting Figure 3: All metabolite correlations for dataset 2 (SPECIAL, 3T). Solely for display purposes, ratios to unsuppressed water and total creatine (Cr+PCr) are normalised to the maximum value fitted by either FSL-MRS or LCModel.

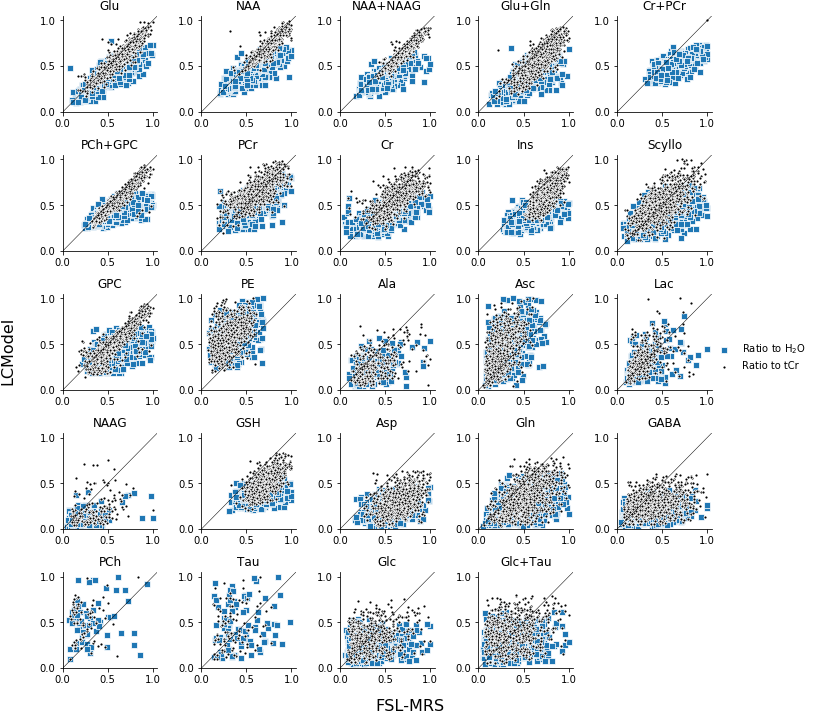

Supporting Figure 4: All metabolite correlations for dataset 3 (MRSI, 3T). Solely for display purposes, ratios to unsuppressed water and total creatine (Cr+PCr) are normalised to the maximum value fitted by either FSL-MRS or LCModel.

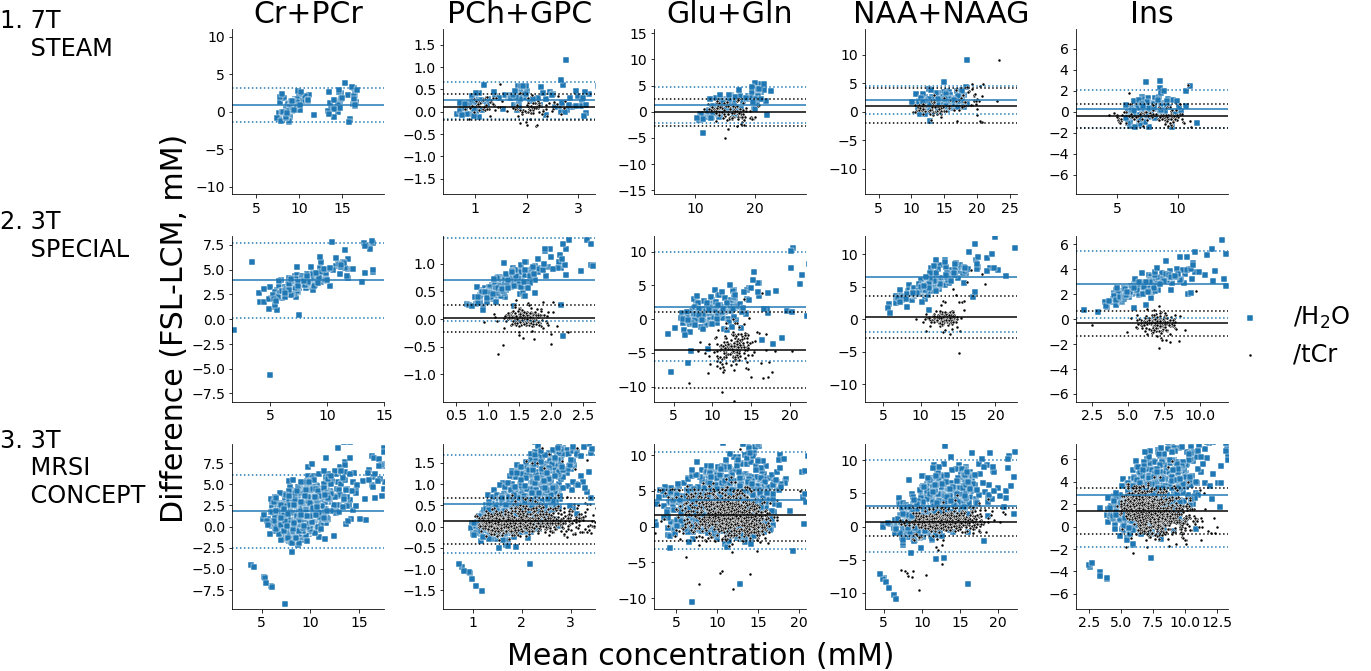

Supporting Figure 5: Bland-Altman plots for selected high-SNR metabolites. Solid horizontal lines mark the bias (mean difference) and dashed lines the limits of agreement (95% CI). Concentrations referenced to total creatine have been multiplied by the mean total creatine concentrations (when referenced to water) to approximate molar concentrations.

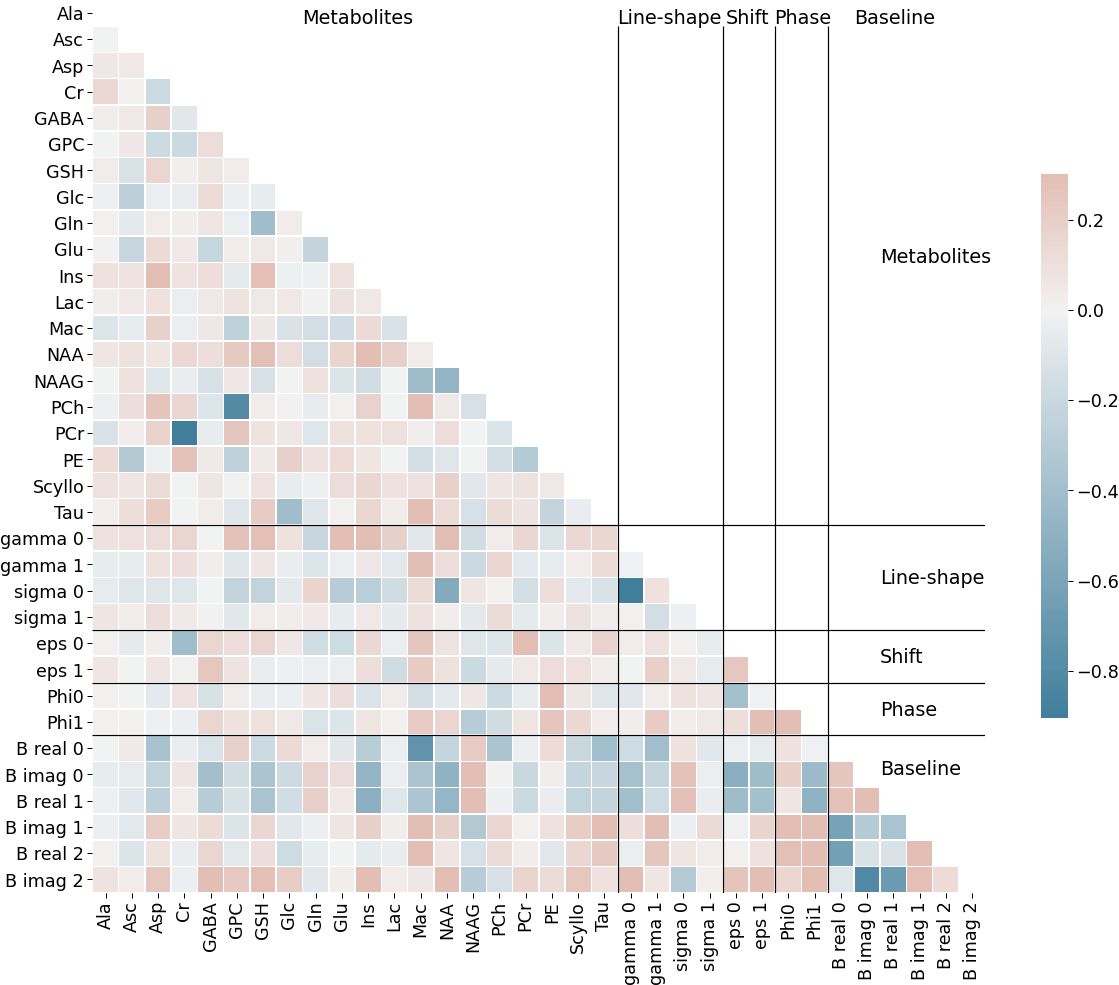

Supporting Figure 6: Correlations between fitting parameters for a single 7T SVS dataset (dataset #1). Fitting was run with 5000 MCMC sampling iterations to ensure convergence.
