## Supplementary material for "FSL-MRS: An end-to-end spectroscopy analysis package": Interactive SVS fitting report

### FSL MRS Report

---

```
    Date        : 2020-06-15 19:11
    FID         : fsl_mrs_preproc/metab.nii
    Basis       : example_data/steam_11ms
    H2O         : fsl_mrs_preproc/wref.nii
```

---

Summary -
Nuisance -
QC -
Uncertainty -
Real/Imag -
Metabs -
Quantification -
Methods

---

### Summary

---

### Voxel location

---

### Nuisance parameters

---

### QC parameters

---

### Uncertainties

---

### Fitting summary (real/imag)

---

### Individual metabolite spectra

---

### Quantification information

|  |  |
| --- | --- |
| Metabolite T_2_: | 160.0 ms |
| Sequence echo time (T_E_): | 11.0 ms |
| T_2_ relaxation corrected water concentration: | 34361 mmol/kg |
| Metabolite relaxation correction (1/e^(-T_E_/T_2_)^): | 1.07 |
| Raw concentration to molarity scaling: | 20.66 |
| Raw concentration to molality scaling: | 27.79 |

---

### Analysis methods

Fitting of the SVS data was performed using a Linear Combination model as described in [1] and as implemented in FSL-MRS version 1.0.0, part of FSL (FMRIB's Software Library, www.fmrib.ox.ac.uk/fsl). Briefly, basis spectra are fitted to the complex-valued spectrum in the frequency domain. The basis spectra are shifted and broadened with parameters fitted to the data and grouped into 2 metabolite groups. A complex polynomial baseline is also concurrrently fitted (order=2). Model fitting was performed using the Metropolis Hastings algorithm.

---
