## Supplementary material for "FSL-MRS: An end-to-end spectroscopy analysis package": Interactive SVS processing report

Report for Combination - 2020-06-15 19:08


### Combined report for fsl\_mrs\_preproc/metab

Combined using merge\_mrs\_reports. Part of the FSL-MRS package.

#### Combination

Report for fsl\_mrs.utils.preproc.combine.combine\_FIDs.
Generated at 19:08:15 on 15/06/2020.

Combination of spectra. Method = svd


#### BadAverageRemoval

Report for fsl\_mrs.utils.preproc.unlike.identifyUnlikeFIDs.
Generated at 19:08:16 on 15/06/2020.

Identification of FIDs unlike others. SD limit = 1.96

#### Align

Report for fsl\_mrs.utils.align.phase\_freq\_align.
Generated at 19:08:21 on 15/06/2020.

Alignment parameters.

Transients before alignment.

Transients after alignment.

#### Combination

Report for fsl\_mrs.utils.preproc.combine.combine\_FIDs.
Generated at 19:08:22 on 15/06/2020.

Combination of spectra. Method = mean

#### ECC

Report for fsl\_mrs.utils.eddycorrect.eddy\_correct.
Generated at 19:08:22 on 15/06/2020.

Eddy correction summary.

Signal + reference phases.

#### HLSVD

Report for fsl\_mrs.utils.preproc.remove.HLSVD.
Generated at 19:08:23 on 15/06/2020.

HLSVD removal of peaks in the range 4.3 to 5.0 ppm.

#### Truncate

Report for fsl\_mrs.utils.preproc.shifting.truncate.
Generated at 19:08:23 on 15/06/2020.

Truncation in time domain.

#### Shift to ref

Report for fsl\_mrs.utils.preproc.shifting.shifttoref.
Generated at 19:08:23 on 15/06/2020.

Frequency shift to reference peak (max in range).

#### Phase correction

Report for fsl\_mrs.utils.preproc.phasing.phaseCorrect.
Generated at 19:08:44 on 15/06/2020.

Phase correction of spectra based on maximum in the range 2.8 to 3.2 ppm.

#### Phase correction

Report for fsl\_mrs.utils.preproc.phasing.phaseCorrect.
Generated at 19:08:44 on 15/06/2020.

Phase correction of spectra based on maximum in the range 4.6 to 4.7 ppm.
